# Genetic meta-analysis of obsessive-compulsive disorder and self-report compulsive symptoms

**DOI:** 10.1101/785311

**Authors:** Dirk J.A. Smit, Danielle Cath, Nuno R. Zilhão, Hill F. Ip, Damiaan Denys, Anouk den Braber, Eco J.C. de Geus, Karin J.H Verweij, Jouke-Jan Hottenga, Dorret I. Boomsma

## Abstract

We investigated whether obsessive compulsive (OC) symptoms from a population-based sample could be analyzed to detect genetic variants influencing OCD. We performed a GWAS on the obsession (rumination and impulsions) and compulsion (checking, washing, and ordering/precision) subscales of an abbreviated version of the Padua Inventory (N=8267 with genome-wide genotyping and phenotyping). The compulsion subscale showed a substantial and significant positive genetic correlation with an OCD case-control GWAS (r_G_=0.61, p=0.017) previously published by the Psychiatric Genomics Consortium (PGC-OCD). The obsession subscale and the total Padua score showed no significant genetic correlations (r_G_=–0.02 and r_G_=0.42, respectively). A meta-analysis of the compulsive symptoms GWAS with the PGC-OCD revealed no genome-wide significant SNPs (combined N=17992, indicating that the power is still low for individual SNP effects). A gene-based association analysis, however, yielded two novel genes (*WDR7* and *ADCK1*). The top 250 genes in the gene-based test also showed significant increase in enrichment for psychiatric and brain-expressed genes. S-Predixcan testing showed that for genes expressed in hippocampus, amygdala, and caudate nucleus significance increased in the meta-analysis with compulsive symptoms compared to the original PGC-OCD GWAS. Thus, inclusion of dimensional symptom data in genome-wide association on clinical case-control GWAS of OCD may be useful to find genes for OCD if the data are based on quantitative indices of compulsive behavior. SNP-level power increases were limited, but aggregate, gene-level analyses showed increased enrichment for brain-expressed genes related to psychiatric disorders, and increased association with gene-expression in brain tissues with known emotional, reward processing, memory, and fear-formation functions.

---

Obsessive-compulsive disorder (OCD) is characterized by recurrent, unwanted thoughts (obsessions) and/or repetitive behaviors (compulsions). The repetitive behaviors or mental acts (such as hand washing, ordering, and checking) are performed in response to an obsession or according to rules that must be applied rigidly. They are aimed at preventing or reducing distress of a feared event or situation, a fear which at the same time is clearly unrealistic and/or excessive. OCD is associated with considerable suffering and markedly impairs individuals’ social and occupational functioning. The lifetime population prevalence of OCD is estimated to be 2 to 3% (Kessler et al., 2005).

Genetic studies have firmly established that OCD has a significant heritable component. A family study has shown evidence for increased odds-ratios in family members of OCD probands (Pauls, Alsobrook, Goodman, Rasmussen, & Leckman, 1995). However, family studies cannot exclude that the shared rearing environment between family members plays a role in the etiology of the disease, thus biasing heritability estimates. Twin studies may overcome this limitation, however, these studies of OCD diagnosis have been limited in sample size (D. S. van Grootheest, Cath, Beekman, & Boomsma, 2005). Twin studies examining obsessive-compulsive symptoms in the general population estimated its heritability to be around 40% (den Braber et al., 2016; D. S. V. Grootheest, Cath, Beekman, & Boomsma, 2007; D. S. van Grootheest et al., 2005; Iervolino, Rijsdijk, Cherkas, Fullana, & Mataix-Cols, 2011; Zilhão et al., 2015). Overall, these studies suggest that a modest, but significant proportion of the liability for OCD is heritable.

OCD is relatively underrepresented in genome-wide association studies (GWAS), both in sample size and number. The largest case-control meta-analysis of GWAS to date included ca. 2800 cases, which falls well short of the >40.000 cases for many other psychiatric disorders (International Obsessive Compulsive Disorder Foundation Genetics Collaborative (IOCDF-GC) and OCD Collaborative Genetics Association Studies (OCGAS) et al., 2018). No significant associations were reported, likely due to the sample size being smaller than required for studies of complex disorders. One solution to increase sample size is to include data from large databases of validated health questionnaires in individuals that have been genotyped, including non-clinical, population based samples with information on obsessive-compulsive symptoms. This requires that the scores on the questionnaire, or the scores on the subscales, reflect the underlying genetic liability and thus genetically correlate with the clinically established diagnosis of OCD. Since very large numbers of participants are a necessity to identify common genetic variants from GWAS, such self-report symptom data may be crucial to increase sample size of existing case-control GWASs. For ADHD, it was recently demonstrated that this is a viable option, provided that the correct statistical meta-analytic technique is used (Demontis et al., 2019). Here, we aim to use a similar approach for a GWAS of OCD. We will first run GWASs on OC symptom scores and subscales from the Padua Inventory and establish whether the genetic variants underlying OC symptoms are associated with those underlying OCD (International Obsessive Compulsive Disorder Foundation Genetics Collaborative (IOCDF-GC) and OCD Collaborative Genetics Association Studies (OCGAS) et al., 2018), by estimating their genetic correlation. Secondly, we will meta-analyze the GWASs on OCD and correlated OC symptom subscales, to examine whether this results in an increase in power to detect underlying genetic variants in a range of follow-up analyses.

## Methods

### Subjects

Twins and their family members (parents, children, siblings) registered at the Netherlands Twin Register (NTR) (Boomsma et al., 2002) were included in the OC symptom analyses. Every two to three years subjects who participate in NTR studies receive selfreport surveys, that contain a variety of questionnaires related to health, personality, demographics, lifestyle and psychiatric disorders. Data on OC symptoms were available for 20528 subjects (N=10285 in year 2002 and N=15803 in year 2005, with N=5560 overlap). Of these, N=8267 (64% female) had genotype data available and were of Dutch ancestry. Their mean age was 41.6 years (SD 15.4; age range between 18–80 years); Supplementary Figure S1 shows the age histogram. Informed consent was obtained from all participants. The study was approved by the Central Ethics Committee on Research Involving Human Subjects of the VU University Medical Centre, Amsterdam, an Institutional Review Board certified by the U.S. Office of Human Research Protections (IRB number IRB00002991 under Federal-wide Assurance - FWA00017598; IRB/institute codes, NTR 03-180).

### Padua-revised (abbreviated) OC symptoms

The OC symptom self-report data were collected by the abbreviated Dutch translation of the Padua Inventory-Revised (Burns, Keortge, Formea, & Sternberger, 1996; Cath, van Grootheest, Willemsen, van Oppen, & Boomsma, 2008; van Oppen, 1992). Supplementary table 1 shows the questions and subscales of the OC symptom scores. The Padua Inventory-revised separates the worry, thought-related items from the behavioral, compulsive items. Six items are included to measure symptoms of impulsive thoughts and rumination. The remaining six items measure behavioral symptoms of OCD, namely checking, washing, and precision (ordering, counting). Since the OC symptoms scale showed strong evidence for skew, we transformed the data with a square-root transformation to minimize the right skew (Zilhão et al., 2015).

### Genotyping and imputation for OC symptoms

A full description of genotyping, pre-imputation QC, and imputation of the NTR OC symptoms GWAS is provided in the supplementary methods. Several critical steps and parameters are presented here. Samples were removed if DNA sex did not match the expected phenotype, if the Plink heterozygosity F statistic was < −0.10 or > 0.10, or if the genotyping call rate was < 0.90. SNPs were removed if the minor allele frequency (MAF) was <0.01, Hardy-Weinberg Equilibrium (HWE) p-value was <1×10-5, call rate was <0.95, or N Mendelian errors was >20. Palindromic AT/GC SNPs with a MAF range between 0.4 and 0.5 were removed to avoid possible strand alignment issues. These were applied to each genotyping platform that was used. After imputation the datasets of each genotyping platform were merged and QC repeated. Ancestry outliers (non-Dutch ancestry) were defined based on Principal Components Analysis (PCA) by projecting 10 PCs from 1000G reference Phase 3v5. We finally filtered on population-based and sample minor allele frequency (MAF) filtered at 0.03. Allele-frequency differences between 1000G reference and sample over 0.20 were removed (10260 SNPs).

### GWAS on OC symptoms

We ran GWASs for the total score on the Padua scale, the obsessions subscale, and the compulsions subscale with ~4.5 M SNPs in a model with Linear Mixed Effects correcting for population stratification and the genetic relatedness between family members using mixed modeling implemented in GCTA (Yang, Lee, Goddard, & Visscher, 2011). Sex, age, age^2^, and 10 population stratification PCs were specified as fixed effects. We created two types of matrices to model genetic relatedness. The first matrix covered the full genetic relatedness matrix including unrelated subjects. The second, family-based matrix was created from the first by setting all relatedness values under 0.05 to zero. This models the additive genetic effects within and between families separately, thus correcting for both family dependence and ancestry dependence in the SNP effects. Both the full genetic relatedness matrix and the family matrix were used in the association analysis. We excluded information of each chromosome out of the relatedness estimations (leave-one-chromosome-out method) so as to avoid adding information from the currently tested SNP in the residual of the linear mixed model. We inspected LD-score regression intercept estimates to test for adequate control of the complex relatedness in the sample.

### Meta-analysis of PGC-OCD-EA and OC symptoms GWAS

We meta-analyzed the results with the PGC-OCD GWAS (International Obsessive Compulsive Disorder Foundation Genetics Collaborative (IOCDF-GC) and OCD Collaborative Genetics Association Studies (OCGAS) et al., 2018) of 2688 cases and 7037 controls of European ancestry. We meta-analyzed this (dichotomous) case-control PGC-OCD GWAS with the (continuous) OC-symptoms GWAS using the genome-wide association meta-analysis (GWAMA) method described in (Demontis et al., 2019) and implemented in R. The population prevalence was set to 0.01 with the actual number of cases and controls entered.

### SNP heritability

SNP heritability for PGC-OCD and OC-compulsions were established with LD-score regression (Bulik-Sullivan, Loh, et al., 2015) to estimate the proportion of variance that could be explained by the aggregated effect of the SNPs. The method is based on the assumption that a regression of the phenotype on the SNP dosage includes effects of all SNPs in LD with the tested SNP. On average, a SNP that tags many other SNPs will have a higher probability of tagging a causal variant than one that tags few other SNPs. Accordingly, for highly polygenic traits, SNPs with a higher average LD-score have stronger effect sizes than SNPs with lower LD-scores. When regressing the effect size obtained from the GWAS against the LD-score for each SNP, the slope of the regression line gives an estimate of the proportion of variance accounted for by all analyzed SNPs. Standard LD-scores were used based on the Hapmap 3 reference panel, restricted to European populations.

### Genetic correlation

We used cross-trait LD-score regression to estimate the genetic covariation between traits based on GWAS summary statistics (Bulik-Sullivan, Finucane, et al., 2015). The genetic covariance is estimated using the slope from the regression of the product of z-scores from two GWAS studies on the LD-score. The estimate obtained from this method represents the genetic correlation between the two traits based on all polygenic effects captured by SNPs. Standard LD-scores were used as provided by Bulik-Sullivan et al. (Bulik-Sullivan, Loh, et al., 2015) based on the 1000 genomes reference set, restricted to European populations.

### Gene-based test and enrichment analysis

We performed MAGMA positional gene based test of association based SNP effects around genes with a 10kbp margin around the 3’ and 5’ UTR as implemented in FUMA (Watanabe, Taskesen, van Bochoven, & Posthuma, 2017). We performed enrichment tests by comparing the top 250 genes from the gene-based tests to several types of annotated gene sets. Ten brain-expression gene sets were selected in FUMA based on the GTEx v7 database. From these tissues, differentially expressed genes (DEGs) were defined as those showed significant up or down regulation compared to the average expression in all other 52 tissues. Enrichment of these genes in the top 250 genes was determined using the hypergeometric test. In addition, we determined whether sets of GWAS catalog reported genes were overrepresented in the top 250 genes. Finally, we compared the significance of these tests between the original PGC-OCD GWAS and the meta-analysis OCD+compulsion symptoms in order to establish whether stronger enrichment could be obtained.

### Expression analysis

To examine to what extent genes associated with OCD+compulsion symptoms meta-analysis are expressed in the brain, we performed S-Predixcan analysis on the meta-analyzed GWAS results. S-Predixcan is based on Predixcan (Gamazon et al., 2015). Predixcan uses RNAseq gene-expression associations present in the GTEx database to build sparse elastic net models, one model for each pair of the 53 tissue and ca. 30,000 genes. Individual SNP data is then used to impute gene expression. These imputed gene-expressions are then associated with the phenotype, resulting in tissue specific associations of gene-expression with the phenotype. S-Predixcan (Barbeira et al., 2016) is an extension of Predixcan and can be used with summary statistics only.

## Results

### GWAS

Supplementary figures S2A-C shows the Manhattan plots for the obsessions, compulsions, and original PGC-OCD GWASs. The respective Q-Q plots are shown in supplementary figures S3A-C.

### Compulsive symptoms show significant genetic correlation with OCD

Table 1 shows the results of the SNP heritability and genetic correlation analysis for the three GWASs using LD-score regression with the original PGC-OCD. The SNP heritability are consistent with previous results (den Braber et al., 2016). The compulsions subscale showed an almost significant heritability at 11.6% (z=1.88, p=0.06). The obsessions subscale showed lower heritability and did not approach significance. The compulsions composite scale (contamination/ordering/counting / checking symptoms) showed a substantial and significant correlation with the PGC-OCD GWAS (r_G_=0.61, p=0.017). The obsessions subscale and the full scale GWAS did not show a significant r_G_ with the PGC-OCD GWAS.

**Table 1.** SNP-based heritability of the Padua inventory full-scale score GWAS, the compulsions and obsessions subscales, and their genetic correlation with the PGC-OCD GWAS.

|  | heritability |  | genetic correlation with PGC-OCD |  |  |  |
| --- | --- | --- | --- | --- | --- | --- |
| | $h^2$ | SE | $r_G$ | SE | z | p |
| compulsions | 0.116 | 0.062 | 0.61 | 0.255 | 2.378 | 0.017 |
| obsessions | 0.058 | 0.065 | -0.02 | 0.322 | -0.068 | 0.946 |
| full scale | 0.102 | 0.060 | 0.42 | 0.254 | 1.672 | 0.095 |

Note. SNP-based heritabilities and genetic correlations were estimated using LD score regression.

### Meta-analysis

Because only the compulsion subscale showed significant rG, this subscale was selected for metaa-nalyzing with the PGC-OCD GWAS. Supplementary Figure S2-D shows the Manhattan plot for the OCD+compulsion symptoms meta-analysis. Figure S3-D shows the associated Q-Q, and figure 1 shows the comparison of the original PGC-OCD Q-Q to the one for the OCD+compulsion symptoms meta-analysis. The inflation median lambdas were comparable for the original and extended GWASs, □ =1.032 and □ =1.033 respectively, indicating a marginally higher lambda for the meta-analysis. Lambdas for the upper 10% percentile were 1.0205 and 1.0405 respectively, indicating a difference in inflation for top SNP effects. LD score regression intercepts were near 1.0 (0.9889, SE=0.0065 and 0.9937, SE=0.0085, for the original PGC-OCD and the OCD+compulsion symptoms meta-analysis respectively), indicating successful control of the ancestry effects.

**Figure 1.**
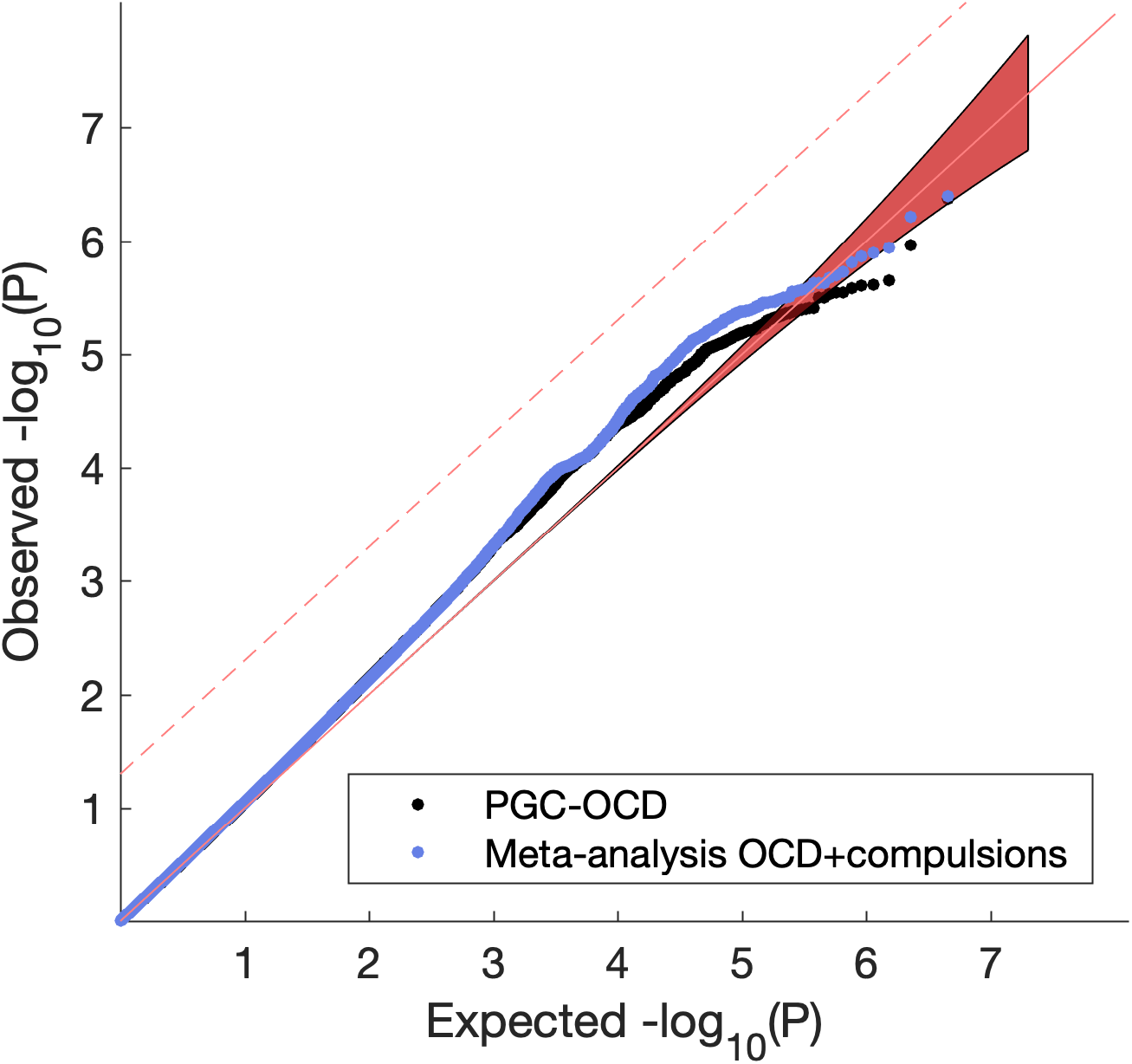
Q-Q plot of observed SNP p-values against expected p-values under the null. Black is the original PGC-OCD GWAS, blue is the meta-analysis of PGC-OCD with compulsions. Dashed line is FDR q=0.05.

### Gene-based tests

Supplementary figure shows the gene-based Manhattan plot, and figure 2 shows the associated Q-Q for the gene-based test results obtained using MAGMA (Leeuw, Mooij, Heskes, & Posthuma, 2015). Four genes were significantly associated with OCD+compulsion symptoms meta-analysis after correcting for multiple testing: *KIT*, *GRID2*, *WDR7* and *ADCK1* at FDR q=0.05. Of these, WDR7 and ADCK1 are novel, whereas the other two were confirm findings from the original PGC-OCD GWAS.

**Figure 2.**
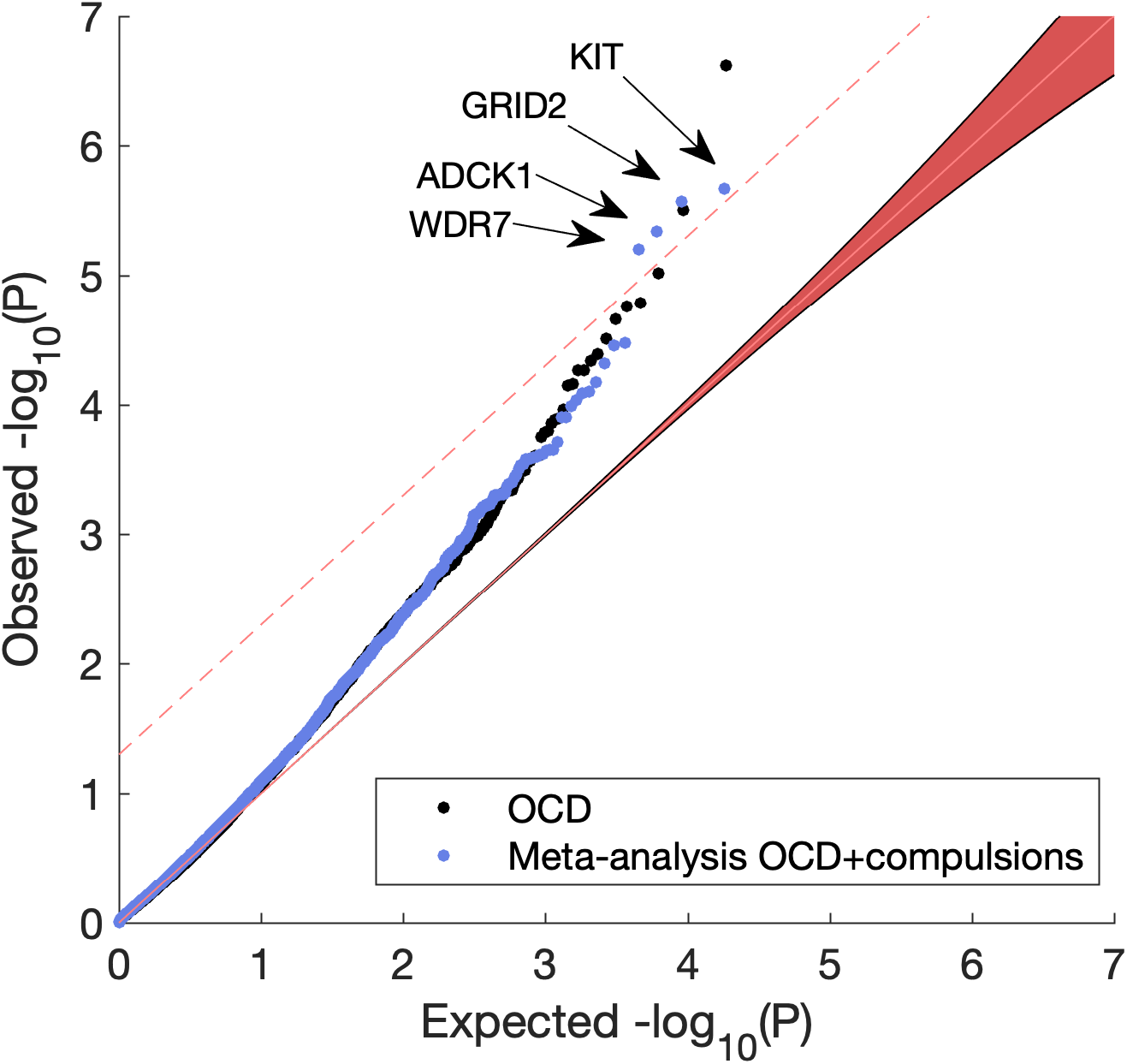
Q-Q plot of observed against expected p-values under the null of the MAGMA gene-based test. Dashed line is FDR q=0.05. The meta-analysis OCD+compulsion symptoms (blue) resulted in four significant discoveries, KIT, GRID2, WDR7, and ADCK1. The latter two are novel findings compared to the original PGC-OCD GWAS (black).

### Altered enrichment of brain expressed genes

We performed an enrichment test of the top 250 genes from the MAGMA positional gene-based analysis, scanned for enrichment of brain-expressed genes form the GTEx database (v7) and for genes reported in GWAS catalog (https://www.ebi.ac.uk/gwas/) associated with psychiatric traits.

Figure 3 shows the significance of enrichment of tissue specific genes in original PGC-OCD before (left; in black) and after meta-analyzing with compulsive symptoms (right, in orange) for all neural tissues excluding cerebellum and spinal cord. Whole blood, and two brain-unrelated tissues (spleen, stomach) were added for reference. Brain-tissue Differentially Expressed Genes (DEGs) were significantly enriched in the PGC-OCD GWAS, with the expressed genes in Anterior Cingulated Cortex (ACC), Nucleus Accumbens (NAcc), and Amygdala as top significant tissues. ACC, Amygdala, frontal cortex, and hippocampus showed strong increases in significance, indicating that more brain-expressed DEGs were present in the top 250 genes of the meta-analysis compared to the original PGC-OCD. Other brain tissues showed marginal change (caudate, putamen, NAcc, hypothalamus). Other tissues (cortex and substantia nigra) showed a decrease in effect.

**Figure 3.**
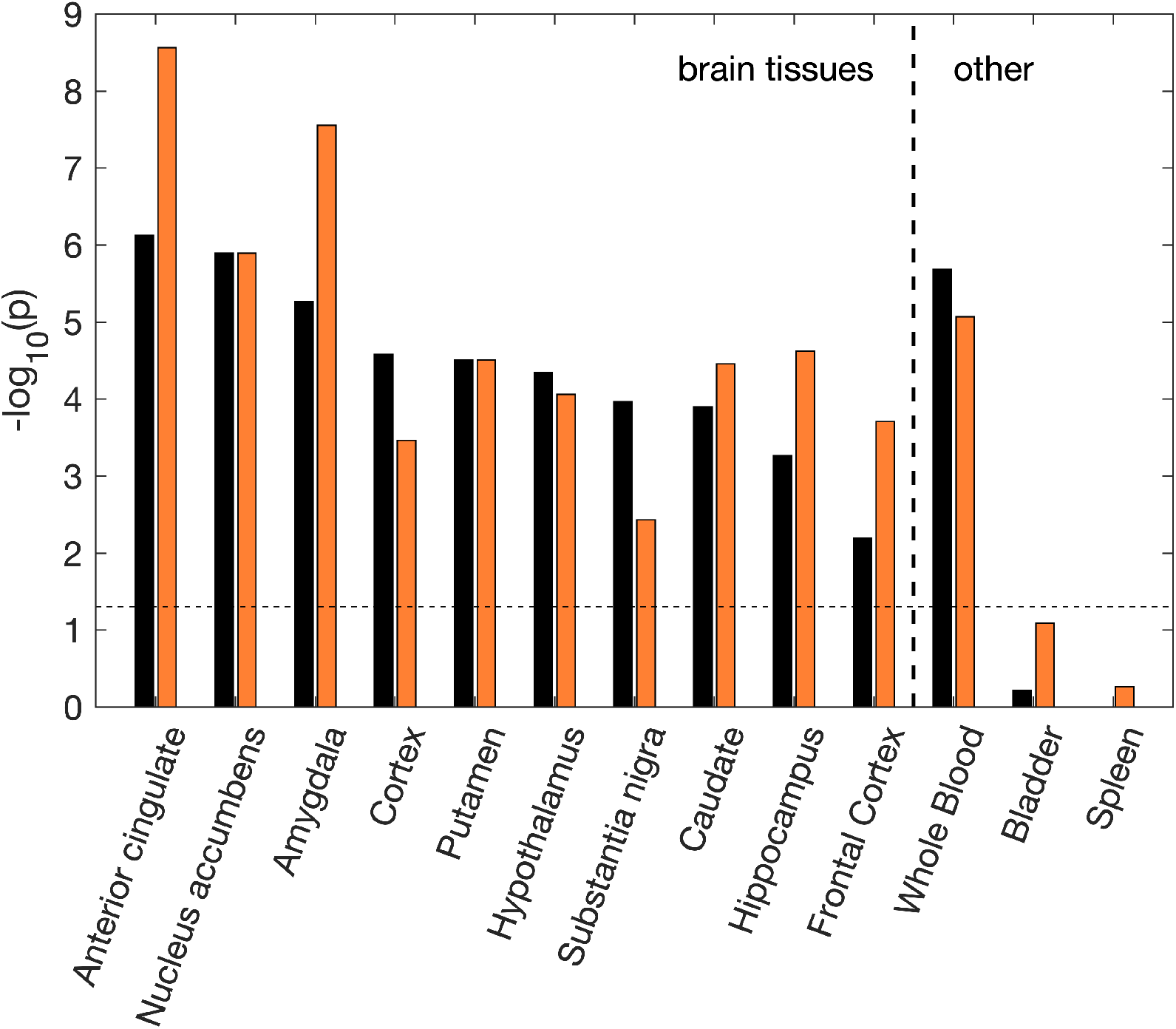
Enrichment results for the PGC-OCD (black) and the OCD+compulsion symptoms meta-analysis (orange). Enrichment of tissue-specific Differentially-Expressed Genes (DEGs) was determined using the hypergeometric test. The y-axis shows the Bonferroni corrected significance of the test as −log10(p). The dashed line indicates the significance threshold (p=0.05). Significance strongly increased for some brain tissues (over 1 point increase for ACC, Amygdala, Hippocampus and Frontal Cortex), and decreased for others (over 1 point decrease for Cortex, Substantia Nigra).

### Enrichment of psychiatric and behavioral gene sets

Gene sets from GWAS catalog are available in FUMA for enrichment analysis. Both the original PGC-OCD GWAS and the meta-analysis with compulsive symptoms showed a large set of significant results after Bonferroni correction, including many psychiatric/behavioral traits. Figure 4 shows the −log10(p) of the enrichment tests of these traits. Schizophrenia genes were strongly enriched, and significance increased over 3 orders of magnitude for the meta-analysis. Significance increased also for “Tourette’s or OCD” and “Autism or Schizophrenia” genes. The schizophrenia genes that contributed to the enrichment effect included the KIT gene, as well as genes on region 3p21 (ITIH4, TMEM112). Genes in this region have been related to bipolar disorder and schizophrenia as well as brain functional activity (Smit et al., 2018).

**Figure 4.**
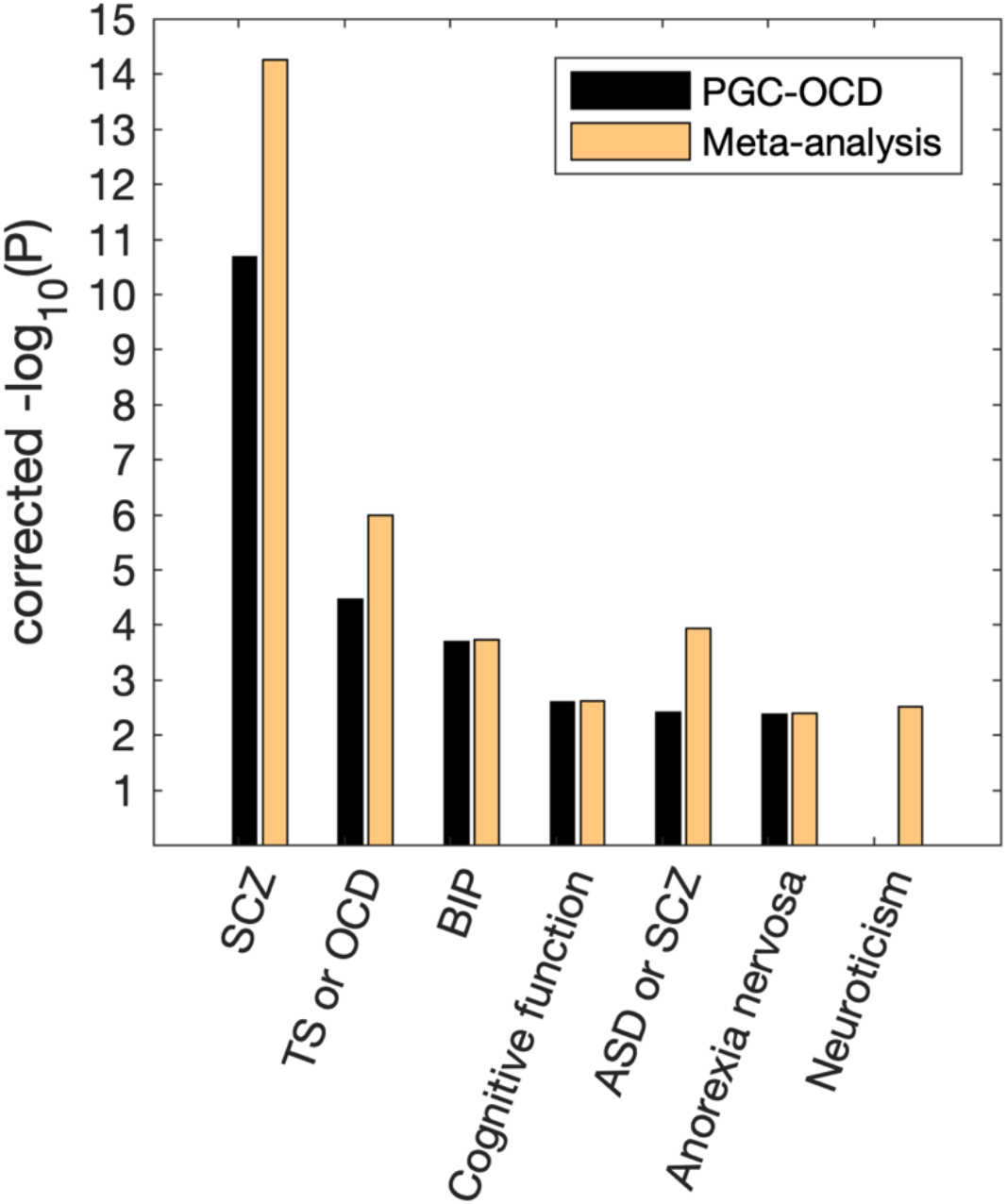
Significance of the enrichment of genes reported in psychiatric/behavioral GWASs, as found in the top 250 genes in the PGC-OCD GWAS and the current meta-analysis. Y-axis shows Bonferroni corrected −log10(p). Either equal significance or a strong increase in the effect was observed when meta-analyzing OCD with compulsive symptoms. SCZ=schizophrenia, TS=Tourette’s syndrome, BIP=bipolar disorder, ASD=Autism spectrum disorders.

### S-Predixcan gene expression associations

Imputed gene expression was not significantly associated with the OCD+compulsion symptoms combined phenotype in any of the ten brain tissues. The expression of RP11-446E9 in Anterior Cingulate Cortex almost reached significance after Bonferroni correction for the number of genes tested within this tissue (p=0.06). Comparing the results from the PGC-OCD GWAS to the meta-analysis, figure 5 shows the difference in log(p-values) for genes that fall under a range of p-value thresholds, with values >0 indicating that the meta-analysis has stronger effects. The figure highlights that increased significance is not present for lower threshold values, but they are for the stronger associations.

**Figure 5.**
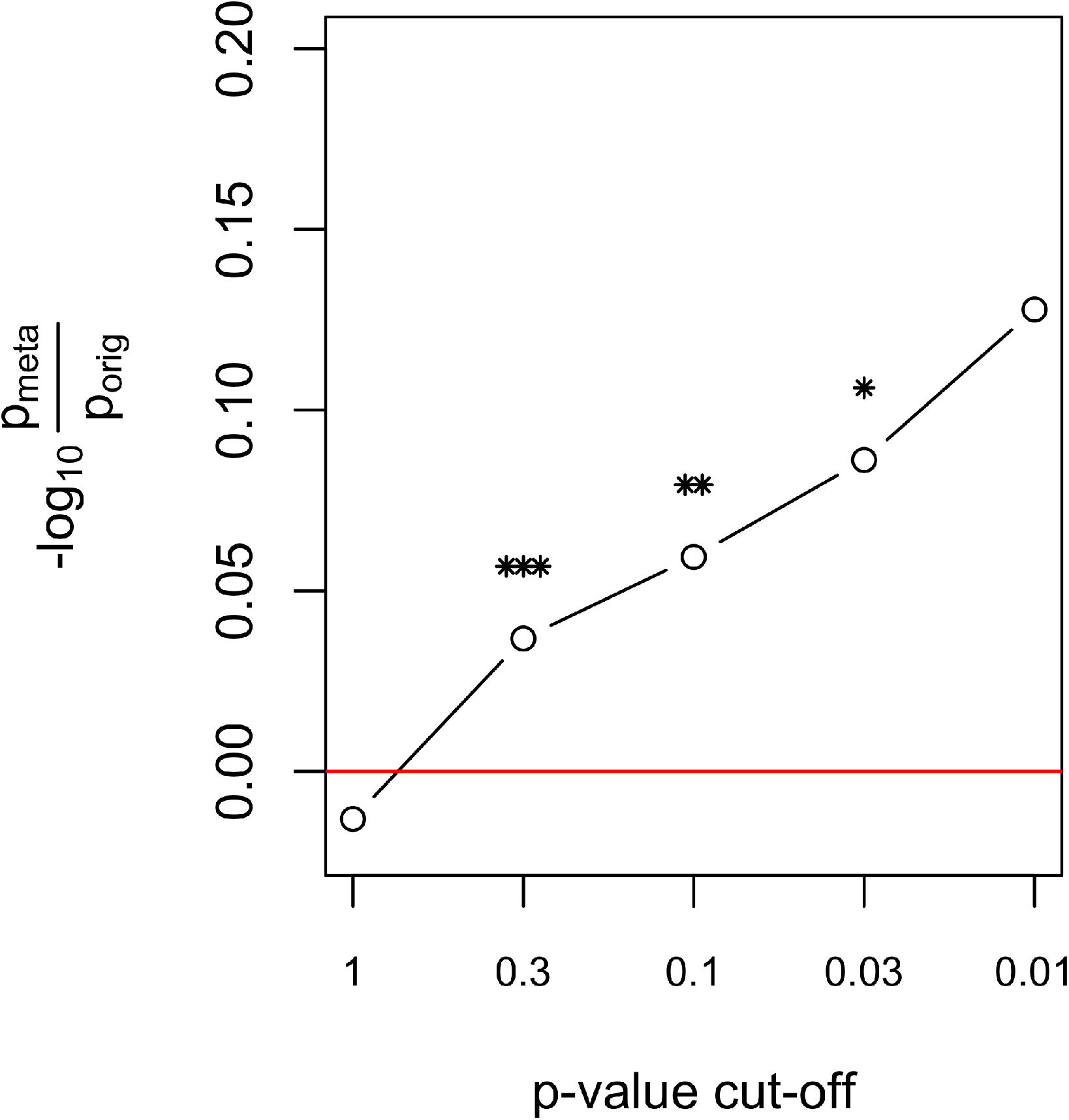
S-Predixcan resulted in stronger p-values after meta-analyzing for the association of a gene’s tissue expression with the phenotype. The figure shows the ratio of the p-values of the meta-analysis to the original GWAS (x-axis), log-transformed. This was performed for genes reaching a specific threshold (x-axis) in the original GWAS. The difference in log p-values for threshold 0.3 to 0.01, indicating stronger effects in the meta-analysis. It also suggests that adding compulsive symptoms only strengthens top genetic expression effects in OCD. The average −log10(p) difference across ten brain tissues is shown. *p<0.05, **p<0.01, ***p<0.001.

## Discussion

We found a substantial and significant genetic correlation between the existing OCD case-control GWAS and our GWAS of compulsion symptoms based on the abbreviated Padua Inventory. For this subscale—which holds questions on behavioral symptoms related to checking behavior, precision (ordering and counting), and contamination fear/washing behavior—the genetic correlation was estimated at r_G_=0.61, p=0.017. The obsessions subscale, on the other hand, was not significantly genetically correlated with the PGC-OCD GWAS. This may reflect the observation that the majority of OCD cases have washing and checking symptoms, thus biasing the OCD GWAS towards these symptoms. The remainder of the Padua Inventory items involve questions on thoughts and worries that may be less specific to OCD. Adding all items scales in equal proportions to the GWAS (i.e. all available Padua item scores) reduced the genetic correlation substantially. Therefore, to optimally add symptom scale analyses to complement case-control GWAS, it might be prudent to include only scores on the questions interrogating the behavioral component of OCD.

Based on the genetic correlation results, we ran the meta-analysis with the PCG-OCD GWAS and the GWAS of compulsive symptoms. This meta-analysis increased the sensitivity of the GWAS to find genes in gene-based and gene-enrichment analyses. We observed two novel discoveries at FDR p=0.05 (ADCK1 and WDR7) in addition to the KIT and GRID2 genes that were previously identified (International Obsessive Compulsive Disorder Foundation Genetics Collaborative (IOCDF-GC) and OCD Collaborative Genetics Association Studies (OCGAS) et al., 2018). SNPs in the WDR7 region have shown trend associations with Tourette’s syndrome/OCD (p=4 x 10^−6^; (Yu et al., 2015)), but note that this report included data from the current GWAS. In addition, SNPs near WDR7 have been suggestively associated with alcohol dependence (p=8 x 10^−6^; (Edwards et al., 2012)). The ADCK1 gene is a novel finding. A SNP near ADCK1 (rs740658277) has been reported in relation to schizophrenia, schizophrenia symptom severity, and response to paliperidone (p=7 x 10^−7^; (Li et al., 2017)). Note that these associations near the WDR7 and ADCK1 genes to psychiatric liabilities are only suggestive, indicating that future research must confirm that these variants are part of the genetic overlap between OCD and other psychiatric disorders.

More substantial power increases were found in the enrichment analysis using gene sets from GTEx and GWAS reports summarized in GWAS catalog. The OCD GWAS showed highly significant enrichment of genes expressed in the Anterior Cingulate Cortex and Nucleus Accumbens. The involvement of these tissues is consistent with their putative role in OCD, reward processing, and as contributing substrates in the cortico-striato-thalamo-cortical circuitry known to be affected in OCD (Denys et al., 2010; Hibar et al., 2018; van den Heuvel et al., 2016). The effects were increased by meta-analyzing the GWAS with the compulsive symptoms GWAS. In addition, the meta-analysis showed increased enrichment of amygdala DEGs, again consistent with the role of this subcortical structure in fear learning and OCD (van den Heuvel et al., 2004).

Gene-set analysis revealed strong increases in GWAS catalog reported genes for traits known to be related to OCD. The original PGC-OCD GWAS showed significant enrichment of genes linked to schizophrenia, consistent with the known genetic overlap between the disorders (Brainstorm Consortium, 2018; Bulik-Sullivan, Finucane, et al., 2015; Martin, Taylor, & Lichtenstein, 2018). This effect increased by over three orders of magnitude in significance after meta-analyzing with the compulsive symptoms GWAS. Likewise, significance of the enrichment of “Tourette’s or OCD” genes increased by over one order of magnitude. The increased enrichment indicates that the top 250 genes were increasingly selective for psychiatric traits known to be genetically correlated with OCD. This indicates that the expression of compulsive behavior is selectively associated with these genes.

The increased enrichments in several psychiatric and brain-expression gene sets was observed without a notable difference in the magnitude of SNP effects between the original OCD GWAS and the meta-analysis. The fact that these power increases were minor after adding compulsive symptoms is likely a consequence of the small size of the compulsion symptoms dataset (N<10k). Even so, the power increases on an SNP-aggregate level observed here suggest that a larger OCD symptoms GWAS could be useful for obtaining increased SNP effects. Moreover, such a power increase is relatively easy to obtain, requiring a relatively short questionnaire with just six compulsion-symptom items from the Padua Inventory.

To summarize, we observed significant SNP-based genetic correlations between the PGC-OCD GWAS and a GWAS of compulsive symptoms in a general population sample. This provided evidence that compulsion symptoms substantially overlap with the genetic liability for clinical diagnosis of OCD, and serves recent calls for doing genome-wide symptom scale analyses to create insight into psychiatric disorder etiology (Davis, 2019). We showed that meta-analyzing the OCD case-control and compulsions symptoms GWAS results has added value in the gene-based and gene-enrichment analyses. This included additional significant genes in the gene-based test and subsequent enrichment analysis of brain expressed genes. These results bode well for larger population-based samples to be merged with clinical samples to increase power for finding the genetic mechanisms underlying OCD.

## ACKNOWLEDGEMENTS

We are indebted to the Psychiatric Genomics Consortium Tourette’s Syndrome / Obsessive-compulsive Disorder workgroup (PGC TS/OCD), in particular Dongmei Yu and Carol Mathews, for making the GWAS summary statistics available. The Genotype-Tissue Expression (GTEx) Project was supported by the Common Fund of the Office of the Director of the National Institutes of Health, and by NCI, NHGRI, NHLBI, NIDA, NIMH, and NINDS. We thank the twins and their family members who participate in the studies of the NTR. This study was supported by the Biobanking and Biomolecular Resources Research Infrastructure BBMRI-NL (NWO 184.021.007 and 184.033.111); FP7-People-2012-ITN, project: TS-EUROTRAIN, grant number 316978; ZonMW (Addiction) 31160008; and European Research Council (ERC-230374); NWO-Groot 480-15-001/674: Netherlands Twin Registry Repository; and Tourette Syndrome Association USA 2008: the genetic epidemiology of tics. Genotyping was realized by grants Rutgers University Cell and DNA Repository (NIMH U24 MH068457-06), the National Institutes of Health (NIH, R01 D0042157-01A1, R01 MH58799-03, MH081802, DA018673, R01 DK092127-04, Grand Opportunity grants 1RC2 MH089951, and 1RC2 MH089995); the Avera Institute for Human Genetics, Sioux Falls, South Dakota (USA). DIB acknowledges the Royal Netherlands Academy of Science Professor Award (PAH/6635). KJHV is supported by the Foundation Volksbond Rotterdam.

## SUPPLEMENTARY INFORMATION

**Table S1.** Padua Inventory (abbreviated) questionnaire items

| Item number | subscale | dimension | question |
| --- | --- | --- | --- |
| 1 | obsessions | impulses | In certain situations, I am afraid of losing my self-control and doing embarrassing things |
| 2 | compulsions | checking | I check and recheck gas and water taps and light switches after turning them off |
| 3 | compulsions | precision | I feel obliged to follow a particular order in dressing, undressing and washing myself |
| 4 | obsessions | impulses | When I see a train approaching I sometimes think I could throw myself under its wheels |
| 5 | compulsions | checking | I return home to check doors, windows , drawers etc, to make sure they are properly shut |
| 6 | obsessions | rumination | When I start thinking of certain things, I become obsessed with them |
| 7 | compulsions | precision | I feel I have to repeat certain numbers for no reason |
| 8 | obsessions | rumination | Unpleasant thoughts come into my mind against my will and I cannot get rid of them |
| 9 | obsessions | rumination | My brain constantly goes its own way and I find it difficult to attend to what is happening round me |
| 10 | compulsions | washing | I sometimes have to wash or clean myself simply because I think I may be dirty or contaminated |
| 11 | obsessions | impulses | I get upset and worried at the sight of knives, daggers and other pointed objects |
| 12 | compulsions | washing | If I touch something which I think is contaminated I immediately have to wash or clean myself |

**Figure S1.**
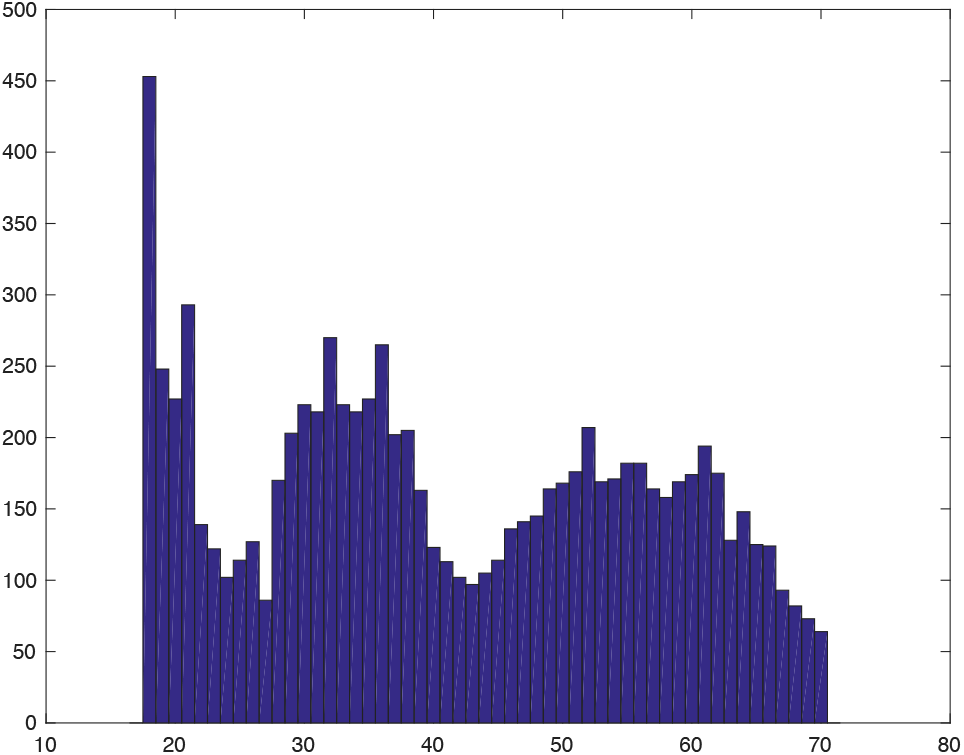
Age distribution histogram.

**Figure S2.**
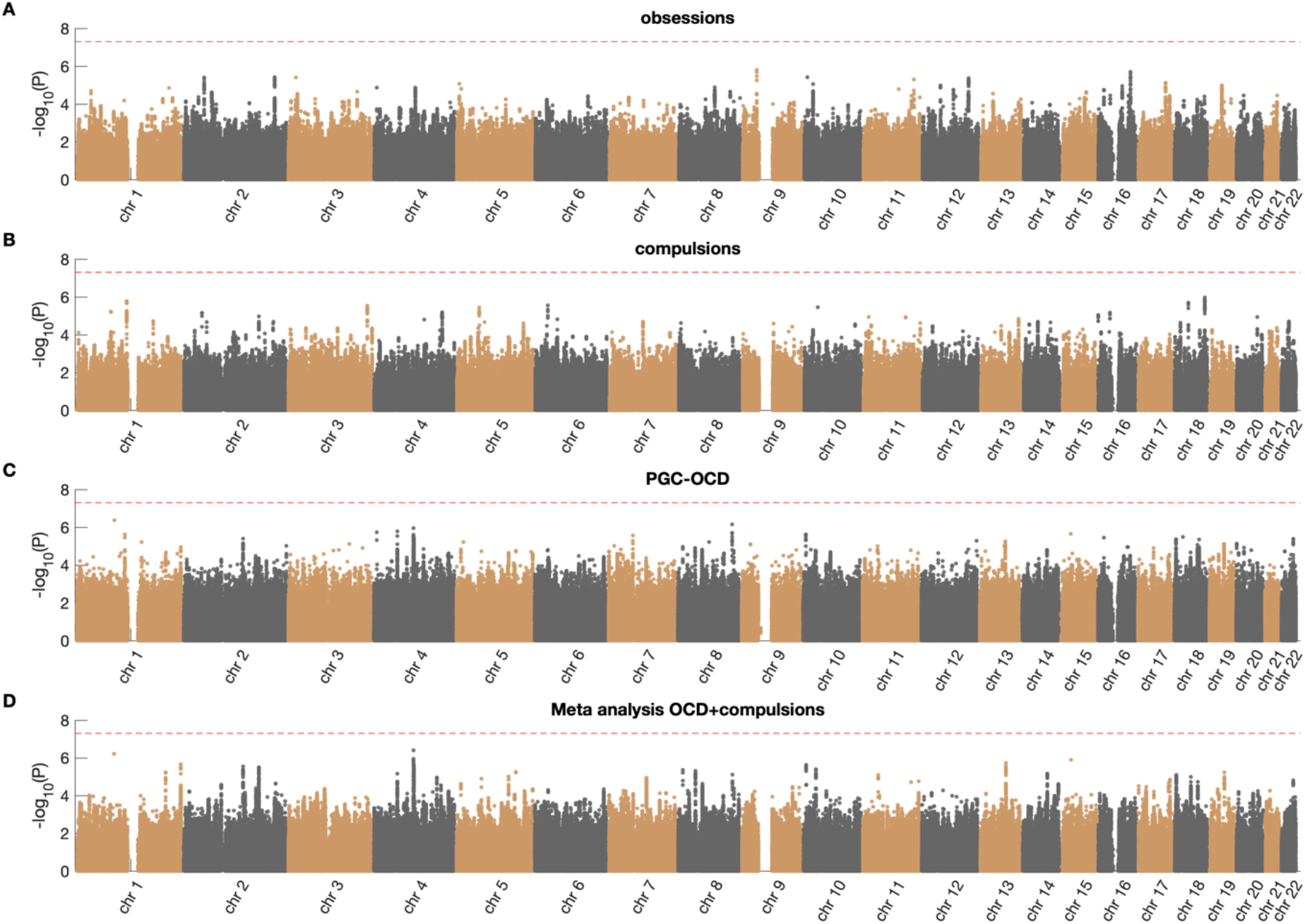
Manhattan plots for the Padua Inventory obsessions subscale (**A**), compulsions subscale (**B**), the original PGC-OCD (**C**), and the meta-analysis of OCD+compulsions (**D**).

**Figure S3.**
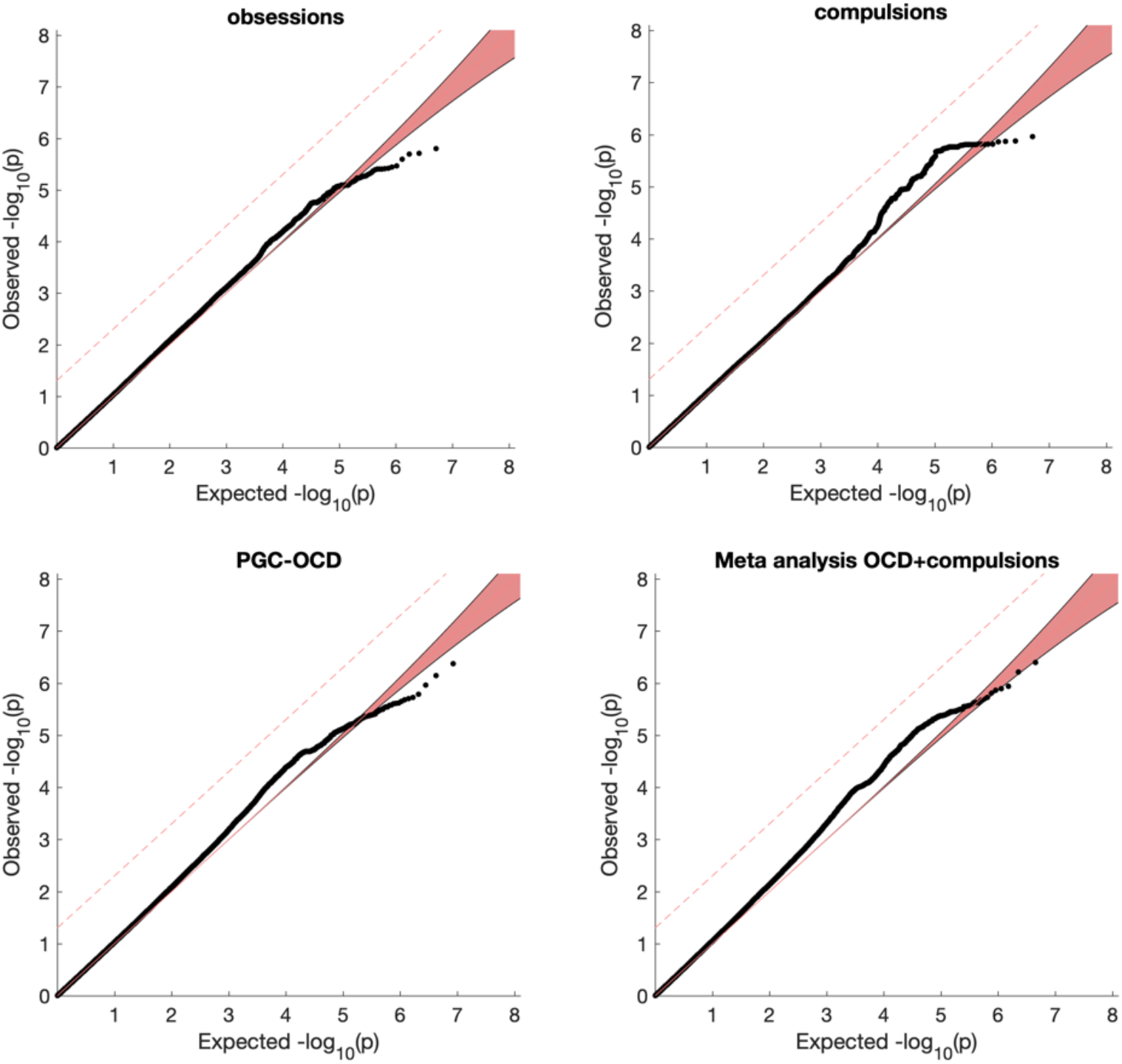
Q-Q plots for the Padua Inventory obsessions subscale (**A**), compulsions subscale (**B**), the original PGC-OCD (**C**), and the meta-analysis of OCD+compulsions (**D**).

**Figure S4.**
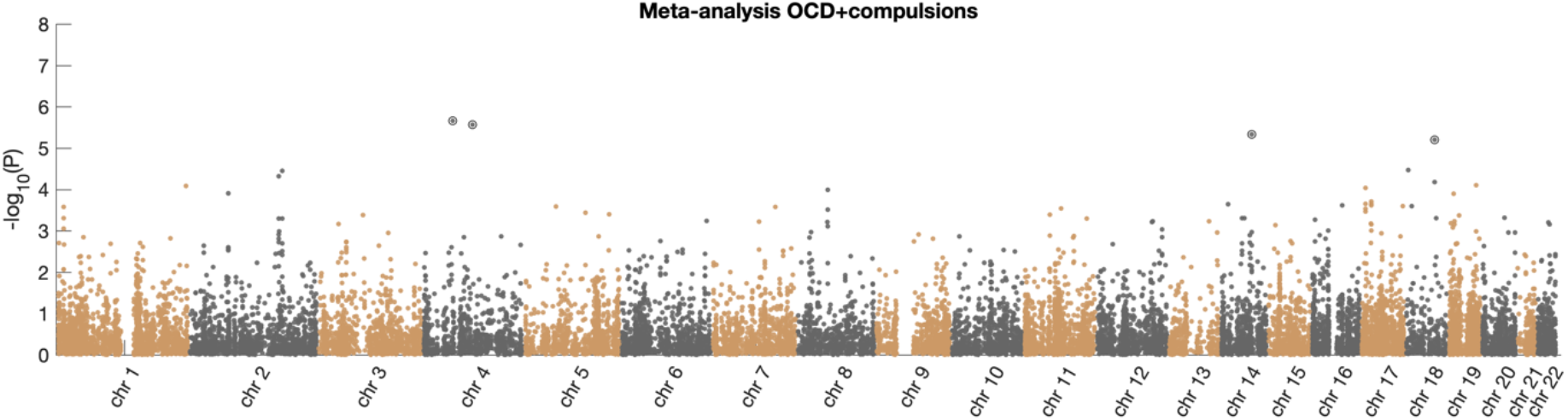
Manhattan plot for the MAGMA gene-based test of association for the meta-analysis OCD+compulsions GWAS.

## Supplementary Methods Netherlands twin registry (NTR): Genetics Methods

### Genotyping, imputation, principal components, GRM analysis

Genotyping was done on multiple platforms over time, namely Perlegen-Affymetrix, Affymetrix 6.0, Affymetrix Axiom, Illumina Human Quad Bead 660, Illumina Omni 1M and Illumina GSA. On each platform genotyping was performed following manufacturers protocols, using the then appropriate calling software. The SNPs of the Perlegen-Affymetrix, Illumina Human Quad Bead 660 and Illumina Omni 1M arrays, which were originally typed on older genome builds, were lifted over to Build 37 HG19 based on RSid locations of the DBSNP 142 marker map. For each genotype platform, samples were removed if DNA sex did not match the expected phenotype, if the Plink heterozygosity F statistic was < −0.10 or > 0.10, or if the genotyping call rate was < 0.90. SNPs were removed if the minor allele frequency (MAF) <0.01, if the Hardy-Weinberg Equilibrium (HWE) p-value < 1×10^−5^, if call rate < 0.95, or if the N Mendel errors > 20 [**#1**]. The absolute value of 20 is used here to remove only the worst offending SNPs (N=2902) in platforms that have familial data present (later this is refiltered more stringently). In addition, palindromic AT/GC SNPs with a MAF range between 0.4 and 0.5 were removed to avoid possible strand alignment issues. For each platform, the data was then position - and strand aligned with the GONL reference set V4. SNPs that had a difference in allele frequency > 0.10, or had mismatching alleles with this reference panel were removed in this step.

The data of the 6 platforms was merged into a single dataset keeping all QCed SNPs of each platform (N=1.781.526). For each individual only one platform was chosen in the following order: Axiom (3144) > Affy6 (8640) > 1M (238) > 660 (1439) > GSA (5938) > Affy-Perl (1238). Based on the ~10.6k SNPs that all platforms have in common, DNA IBD was estimated for all individual pairs using the PLINK and KING programs [**#1,#2**]. These estimates were then compared to the expected familial relations, and samples were removed if these failed to fit. A similar approach was used for DNA zygosity mismatches. Duplicate monozygotic twins N=3032, triplets N=7 as well as NTR samples present in the GONL data N=364 (plus their MZ-twins) were removed from the data prior to imputation. The data were then cross-platform phased and imputed using MACH-ADMIX to predict the missing SNP genotypes in each platform as compared to the other platforms, based on the complete GONL reference panel haplotypes for the SNPs that were present in at least one platform (the ~1.78m) [**#3–#6**]. Post imputation, the 2nd (and 3rd) MZ, plus the GONL samples were re-added duplicating the data from the 1st imputed MZ twin, and the complete SNP data from the GONL reference panel.

After this imputation, SNP QC was redone, now using the full merged dataset with all missing genotypes imputed. SNPs were removed if the HWE p-value was < 1×10^−5^, Mendel error rate was more than mean + 3sd, the R^2^ imputation quality metric was < 0.90, if p-value for association with a single platform vs. all others was < 1×10^−5^. No MAF filter was re-applied (min = 0.0025). This left a cleaned merged dataset of 21.001 NTR individuals with 3032 MZ pairs, 7 MZ trios, and 1.314.639 SNP markers (NchrX=20.792). This cross-platform imputed set described is what we consider to be our ‘genotyped’ dataset in the next two steps, which are the detection of ethnic outliers and the imputation to the 1000 genomes Phase 3v5 and the Human Reference Consortium (HRC) panels [**#7,#8**].

Ancestry outliers (non-Dutch ancestry) were defined based on Principal Components Analysis (PCA) by projecting 10 PCs from 1000G reference set populations on the NTR cross-platform imputed data using the SMARTPCA program as described earlier [**#9,#10**]. Individuals with PC values located outside of the range of European and/or British populations were defined as outliers (N=1823). Upon exclusion of outliers, 10 PCs were recomputed for NTR cross-platform imputed data to capture the variation within the Netherlands.

Genotype imputation to the HRC 1.1 (~40m SNPs) and 1000G Phase 3 version 5 (~49m SNPs) reference panels was done on the Michigan Imputation server on the cross-platform imputed data [**#11**]. For each reference panel the data were aligned using the PERL based “HRC or 1000G Imputation preparation and checking” tool v4.2.5 [**linkonly#12**]. The remaining SNPs (1.302.481 : 1000G, 1.307.940 : HRC) were then phased with EAGLE for the autosomes, and SHAPEIT for chromosome X and then imputed using Minimach 3 following the standard imputation procedures of the server [**#13,#14**].

The cross-chip imputed data was filtered for SNPs having a MAF < 0.01. Then Genetic Relationship Matrices (GRM) were computed for each individual chromosome (1 to 22) using the GCTA software [**#15**]. Subsequently, these 22 matrices were merged into single autosomal matrix. Leave One Chromosome Out (LOCO) GRMs in order to control for genetic background when running genome wide associations, were calculated likewise by merging 22 subsets of 21 chromosomes respectively. Finally, a specific family based GRM was made by setting all individual pairs sharing less than 0.05 of their genome to 0 in the matrix and leaving the other pairs as calculated [**#16**]. No GRMs were calculated for the imputed data (100G and HRC) as these are sufficient to control for confounding in GWAS and to calculate the heritability for various traits.

